# Dynamic polarization of heparan sulfate proteoglycans is involved in planar cell polarity and localization of endogenous Wnt11 during vertebrate neural tube formation

**DOI:** 10.1101/2025.08.12.669988

**Authors:** Minako Suzuki, Ritsuko Takada, Tomoe Kobayashi, Makoto Matsuyama, Shinji Takada, Yusuke Mii

## Abstract

Planar cell polarity (PCP) on a tissue plane is established by asymmetric polarization of core PCP components in individual cells. Recently, we found that Wnt11 organizes PCP in *Xenopus* embryos by accumulating locally with core PCP components to be polarized, but not by acting as a directional cue forming a global gradient. Since heparan sulfate proteoglycans (HSPGs) associate with Wnt on cell membranes, how HSPGs are involved in Wnt polarization and PCP formation is a central issue. Here, we show that HSPGs are dynamically organized in a polarized manner, as with core PCP components and Wnt11, by associating with them. While HSPGs with HS chains enriched by *N*-sulfation and *N*-deacetylation are polarized, depending on Wnt and core PCP components, NDST1, which catalyzes these modifications, is required for Wnt11 polarization and PCP formation. Thus, mutual regulation among HS chains, Wnt11, and PCP components appears to be essential for PCP formation.

**Highlights:**

- *ndst1* is required for Wnt11 localization and proper PCP formation.
- NDST1-modified HS chains are polarized in the neural plate during development.
- Polarization of NDST1-modified HS chains depends on PCP formation.
- Positive feedback loop results in polarized distribution of HSPGs, Wnt11, and PCP.

**Graphical abstract:** 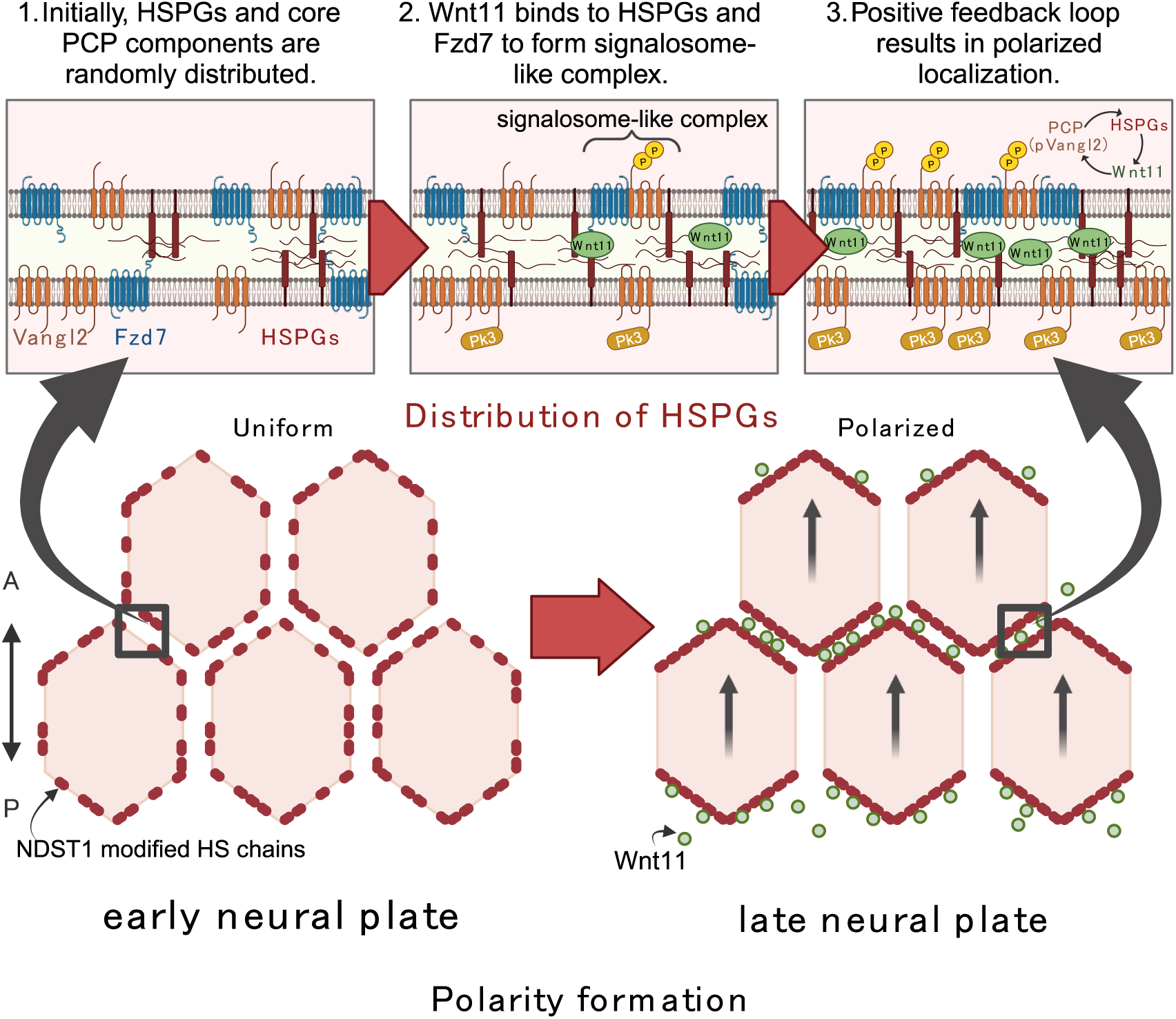

## Introduction

Planar cell polarity (PCP) refers to coordinated alignment of cell orientations in a tissue plane. Along this polarization, cell shape and behavior and tissue morphology are organized by modulating cell-cell adhesions and cytoskeletons ^1–5^. PCP is established by a group of essential proteins known as core PCP components, including Frizzled (Fzd), Dishevelled (Dvl), Van Gogh-like (Vangl; Vang aka Strabismus, Stbm in *Drosophila*), Prikkle (Pk), and Celsr (Flamingo aka Starry Night in *Drosophila*). These core PCP components are thought to form complexes, localizing asymmetrically on cell membranes perpendicular to the polarity axis ^1–5^. For example, in the neural plate of vertebrate embryos, where PCP is established along the antero-posterior (AP) axis, Pk and Dvl are enriched on the anterior and posterior edges of each cell, respectively^2,6^.

Wnt has been considered a morphogen, which generally forms a concentration gradient to provide positional or directional information to cells in tissues. In terms of PCP, noncanonical Wnt signaling (β-catenin-independent signaling) has been considered a possible source of global cues that coordinate polarity of cells on a tissue-wide scale. Consistent with this idea, ectopically expressed Wnt can direct PCP in many cases, including *Drosophila* wing, ectoderm of early *Xenopus* embryos, and the node of mouse embryos ^7–9^. In contrast, recent studies have demonstrated that Wnt ligand is dispensable for normal PCP formation in *Drosophila* ^10,11^. In vertebrates, the requirement of Wnt for PCP is widely accepted as supported by loss-of-function studies with Wnt5a and/or Wnt11, and of downstream noncanonical Wnt signaling, all of which result in defects of axial elongation, neural tube closure, and/or convergent extension movements (CEMs) ^12–14^. However, it is still debatable whether a gradient of Wnt ligands is required for PCP, and whether Wnt ligands serve instructive or permissive functions in regard to PCP. Moreover, mechanisms by which Wnt polarizes PCP, or establishes asymmetry of core PCP components, remain largely unknown. On the other hand, our recent study showed that Wnt11 does not form a concentration gradient in *Xenopus* neural plate ^15^ (Figure 1A left), and is polarized like core PCP components (Figure 1A right). However, that study also proposed that Wnt11 does not serve as the primary global cue for PCP, but that local interaction between Wnt11 and core PCP components can establish PCP in a self-organizing manner. ^15^.

**Figure 1.**
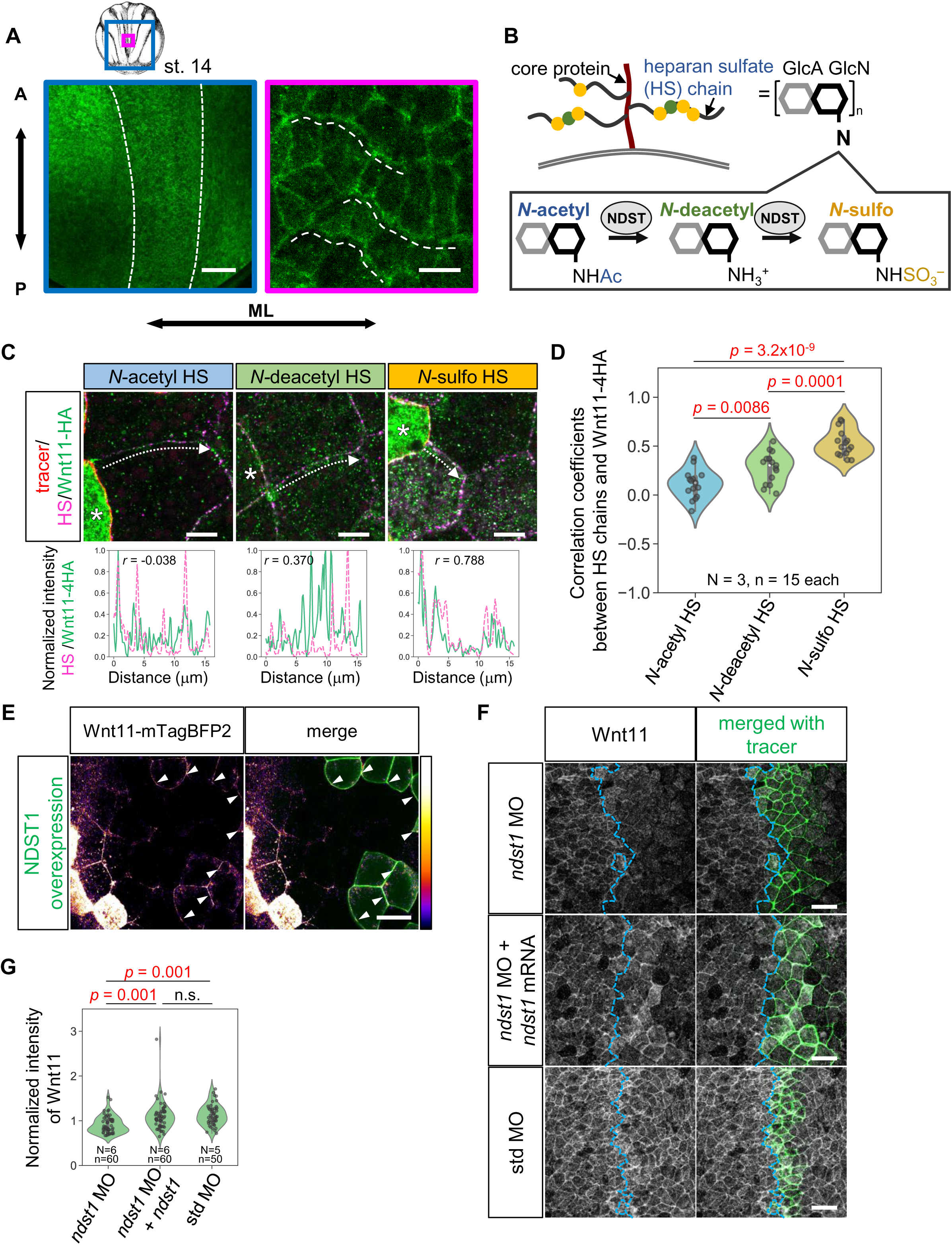
NDST1-modified HS chains are required for endogenous Wnt11 localization. (A) Schematic image of neural plate Stage 14 *Xenopus* embryo and endogenous Wnt11 distribution in the *Xenopus* neural plate. White dashed lines indicate medio-lateral (ML) localization of Wnt11. (B) Schematic image of HSPGs and modification of HS chains by NDST1. (C) Co-localization of Wnt11-4HA and each modified state of HS chains. Fluorescent intensity at cell boundaries along white dotted arrows was plotted. Asterisks show Wnt11-4HA expressing cells. Fluorescent intensities were normalized with Min-Max normalization in each plot. (D) Average of correlation coefficients between Wnt11-4HA and HS chains at cell boundaries. Statistical analysis was performed with the Shapiro-Wilk test to assess data normality and was followed by Tukey’s HSD test for multiple comparisons. Numbers of embryos (N) and numbers of cell boundaries (n) are as indicated. (E) Wnt11-mTagBFP2 accumulation on NDST1-overexpressing cells. Live-imaging at Stage 14. mRNAs of *wnt11-mTagBFP2* and a membrane tracer (mEGFP-Kras) and mRNAs of *ndst1* and another membrane tracer (mRuby2-Kras) were injected into different ventral blastomeres of 4-cell stage embryos. Arrowheads indicate accumulation of Wnt11-mTag-BFP2. The lookup table used for Wnt11-mTagBFP2 is indicated on the right. (F) Requirement of *ndst1* for membrane localization of endogenous Wnt11 in the neural plate (Stage 14). MOs and mRNAs of membrane tracer (mRuby2-KRas) and *ndst1* were co-injected into the right dorsal blastomere of 4-cell stage embryos, targeting the future neural plate. Boundaries of tracer-negative and -positive area are indicated with dashed lines. (G) Quantification of Wnt11 membrane localization in *ndst1* knockdown experiments (F). Statistical analysis was performed with the Shapiro-Wilk test to assess data normality and was followed by the Kruskal-Wallis test for pre-analysis and the Steel-Dwass-Critchlow-Fligner test for multiple comparisons. Numbers of embryos (N) and numbers of cell boundaries (n) are as indicated. Amounts of mRNAs/MOs: *wnt11-mTagBFP2,* 1.0 ng/embryo (E); *mRuby2-kras,* 100 pg/embryo (E) or 67 pg/embryo (F); *mEGFP-kras,* 100 pg/embryo (E); *ndst1,* 100 pg/embryo (E) or 33 pg/embryo (F and G); *ndst1* MOs and std MO, 21 ng/embryo (F and G). Scale bars, 200 μm (A, left), 20 μm (A, right), 10 μm (C), 25 μm (E), 50 μm (F).

Heparan sulfate proteoglycans (HSPGs) consist of core proteins, such as Glypican of GPI-anchored protein, Syndecan of transmembrane protein, and heparan sulfate (HS) glycosaminoglycan chains attached to core proteins ^16^ (Figure 1B). HS chains are synthesized as repeats of disaccharide units (typically 40-160 units) of glucuronic acid and *N-*acetylglucosamine (GlcA-GlcNAc) by co-polymerase Ext. Modifications of HS chains are initialized by *N-* deacetylation and subsequent *N*-sulfation of GlcNAc by *N*-deacetylase/*N*-sulfotransferase (NDST, *sulfateless* mutant in *Drosophila*) (Figure 1B), followed by epimerization of GlcA to iduronic acid (IdoA) and/or *O*-sulfations ^16^. HSPGs serve as essential scaffolds for extracellular distribution of morphogens, including Wnt ^17^. For instance, *Drosophila* genetics have identified a number of HSPG-related genes that are required for proper distribution and function of Wingless (Wg, Wnt1 ortholog in *Drosophila*) ^17–19^. HSPGs are also involved in noncanonical Wnt signaling and PCP formation. Both loss and gain of function mutations of glypican4 (*gpc4*, *knypek* mutant in zebrafish) result in defects of axial elongation and CEMs, similar to disruption of noncanonical Wnt signaling ^20,21^, and phenocopying disruption of noncanonical Wnt signaling ^12^. Indeed, *knypek*/*gpc4* interacts genetically with *wnt11*, suggesting their functional association ^21^. Gpc4 also physically interacts with Wnt11 in *Xenopus* embryos ^22^. Syndecan4 (Sdc4) is thought to be involved in morphogenesis of *Xenopus* embryos, probably by modulating noncanonical Wnt signaling ^23^. As for PCP, involvement of HSPGs has also been shown. Indeed, genetic interaction between *vangl2* and *gpc4* or *sdc4* have been reported ^20,24^. Enzymatic digestion of HS chains causes an open neural tube phenotype in wild-type mice, and this phenotype is enhanced in a heterozygous background of the dominant negative *Vangl2* allele, *Loop-tail* ^25^. Thus, HSPGs, especially HS chains and their modifications, are important for noncanonical Wnt signaling and PCP formation. However, how HSPGs are involved in regulation of PCP and noncanonical Wnt signaling is largely unknown.

Although HSPGs have been intensively studied in genetic and biochemical analyses, cell biological understanding of HSPGs still remains largely unknown. By visualizing HS chains using specific monoclonal antibodies ^26–28^, we demonstrated that HSPGs form individual clusters that differ in HS chain modifications, especially, *N-*acetylation-rich HS (*N-*acetyl HS) clusters and *N-* sulfation-rich HS (*N-*sulfo HS) clusters on cell surfaces in *Xenopus* embryos. Furthermore, *N-* sulfo HS clusters are important for spatial distribution and signal transduction of Wnt8 ^29^.

Differences in these modifications of HS chains may be a key to understand detailed functions of HSPGs, including their function in PCP. Interestingly, modifications of HS chains appear to be required for PCP because *NDST1* knockout mice show various morphological defects including an open neural tube, which is also a common phenotype in PCP defects, although relationships between NDST and PCP have never been mentioned ^30^. However, it is still unclear how modifications of HS chains are involved in establishment of the polarized pattern of Wnt11 and core PCP components.

In this study, to investigate involvement of HS modification in PCP formation and the polarized distribution of Wnt11, we visualized HS chains with monoclonal antibodies specific to modification states of HS chains, especially *N*-acetylation, *N*-deacetylation, or *N*-sulfation. Moreover, we manipulated these HS modifications by loss-of- or gain-of-function of *ndst1.* Surprisingly, we found that spatial distribution of HSPGs was not static, but dynamic, forming polarized patterns in *Xenopus* neural plate. We also demonstrated that this dynamic change of HSPG distribution is likely achieved by association with core PCP components and Wnt11. Our findings provide mechanistic insights into relationships among HSPGs, PCP, and noncanonical Wnt signaling, and further reveal a novel concept of dynamic pattern formation of HSPGs, changing our view of HSPGs as static and uniform.

## Results

### Exogenous Wnt11 colocalizes with NDST1-modified HS chains, and with *N*-deacetyl and *N*-sulfo HS

We previously proposed that distribution of secreted signal proteins in tissues or embryos is regulated by interaction with scaffold molecules on cell membranes ^31^. In particular, HSPGs enriched with *N-*sulfo or *N*-acetyl HS, cluster separately on cell membranes, and *N*-sulfo-rich HSPG clusters act as scaffolds to accumulate Wnt8 in *Xenopus* embryos ^29^. First, we asked whether *N-*deacetylated HS, which is also generated by NDST, forms distinct clusters on cell membranes, as well as *N*-acetyl and *N*-sulfo HS. In this study, we used the following monoclonal antibodies: NAH46 for *N-*acetyl HS ^27^, JM-403 for *N*-deacetyl HS ^26^, and HepSS-1 for *N*-sulfo HS ^28,32^. Because all of these antibodies were mouse IgMs, double staining of *N*-deacetyl and *N*-sulfo HS was performed with specific methods to avoid cross-reaction (Figure S1A). Weak correlation between *N*-deacetyl and *N*-sulfo HS was found; however, clusters enriched with *N*-deacetyl or *N*-sulfo HS chains were also found on cell membranes (Figures S1B to S1D), suggesting *N*-deacetyl HS-rich clusters. Based on these findings, we investigated whether these modification states of HS chains co-localize with Wnt11 on cell membranes. To this end, HA-tagged Wnt11 (Wnt11-4HA) was overexpressed in the ventral animal cap region and co-localization between Wnt11-4HA and HS chains was examined. Both Wnt11-4HA and HS chains were visualized with whole-mount immunostaining using anti-HA and three types of anti-HS chain antibodies. Since these antibodies recognize different clusters on cell membranes, it is thought that HSPGs enriched with similar HS modifications are locally clustered on cell membranes. However, HS chains are polymerized as units of GlcA-GlcNAc, and modification of disaccharide units in HS chain is not likely to be uniform. In this paper, clusters of HSPGs stained with these three antibodies are referred to as *N*-sulfo HS, *N*-acetyl HS, and *N*-deacetyl HS, respectively, but this does not mean that all disaccharide units in each cluster are similarly modified. Co-localization was represented with correlation coefficients between Wnt11-4HA and each type of HS chains (Figures 1C and 1D). Compared with *N*-acetyl HS, localization of *N*-deacetyl and *N*-sulfo HS exhibited significantly higher correlation with that of Wnt11-4HA. In particular, *N-*sulfo HS exhibited the highest correlation with Wnt11-4HA among the three types of modified HS chains. This specificity to HS is similar to the case of Wnt8 ^29^.

Because *N*-sulfation and *N*-deacetylation of HS chains, which were well co-localized with Wnt11-4HA, are modified by NDST1, we investigated whether NDST1 overexpression promotes localization of Wnt11 on cell membranes. NDST1 and mTagBFP2-tagged Wnt11 (Wnt11-mTagBFP2) ^15^ were overexpressed in different cells in the ventral animal region of *Xenopus* embryos, and the effect of NDST1 on distribution of Wnt11-mTagBFP2 was examined. Wnt11-mTagBFP2 that diffused from its source cells was strongly accumulated on NDST1 overexpressing cells (Figure 1E, arrowheads), suggesting that HS chain modifications by NDST1 promotes association with Wnt11. From these data, HSPGs clusters enriched with *N*-deacetyl and *N*-sulfo HS can be scaffolds for accumulating Wnt11 on cell surfaces.

### NDST1 is required for polarized localization of endogenous Wnt11 in the neural plate

We next asked whether HS modifications by NDST1 are necessary for the characteristic localization of Wnt11 in the neural plate. *ndst1* was knocked down by injection of antisense morpholine oligos (MO) ^29^ to a dorsal blastomere of 4-cell stage embryos, targeting the right half of the neural plate at Stage 14. *N*-sulfation- and *N*-deacetylation-rich HS chains were decreased by the *ndst1* MO in the neural plate at Stage 14, showing that *ndst1* MO specifically inhibited the function of NDST1, as expected (Figures S2A and S2B). We performed whole-mount immunostaining for endogenous Wnt11 and measured Wnt11 intensity along cell boundaries. Localization of endogenous Wnt11 on cell surfaces was significantly reduced in *ndst1* MO-injected regions compared to standard (std) MO injected ones (Figures 1F and 1G). The effect of the *ndst1* MO was partially rescued by addition of *ndst1* mRNA, showing that the effect of *ndst1* MO was specific (Figures 1F and 1G). These data indicate that *ndst1* is required for membrane localization of endogenous Wnt11 in the neural plate.

### NDST1-modified HS chains are required for proper PCP formation

Since impaired function of Wnt11 could lead to PCP defects ^8,12,33^, we examined morphological abnormalities caused by knockdown of *ndst1*. For this purpose, *ndst1* MO was injected into both dorsal blastomeres in 4-cell stage embryos, targeting the future neural plate. At Stages 21-22, when the suture of the neural tube is completely closed in normal development ^34^, knockdown of *ndst1* caused neural tube closure defects, which can be rescued by *ndst1* mRNA (Figures 2A and 2B; see also Figures S3A and S3B for later stage). Since neural tube closure defects are typically observed with disruption of PCP or noncanonical Wnt signaling ^1,6,35,36^, we further examined whether knockdown of *ndst1* results in disruption of PCP by monitoring localization of core PCP components. In *Xenopus* neural plate, localization of Vangl2, one of the core PCP components, is polarized ^6,37^. We found that the amount of Vangl2 localized on cell membranes was significantly reduced by *ndst1* knockdown in the right half of the neural plate at Stage 14 (Figures 2C and 2D). This reduction was partially rescued by *ndst1* mRNA (Figures 2C and 2D). These data suggest that NDST1 is required for proper PCP formation, in addition to Wnt11 localization, in *Xenopus* embryos. Consistently, overexpression of NDST1 promoted membrane localization of Vangl2 (Figures 2E and 2F). Thus, NDST1-mediated modifications of HS chains are necessary and sufficient for proper PCP formation in the neural plate.

**Figure 2.**
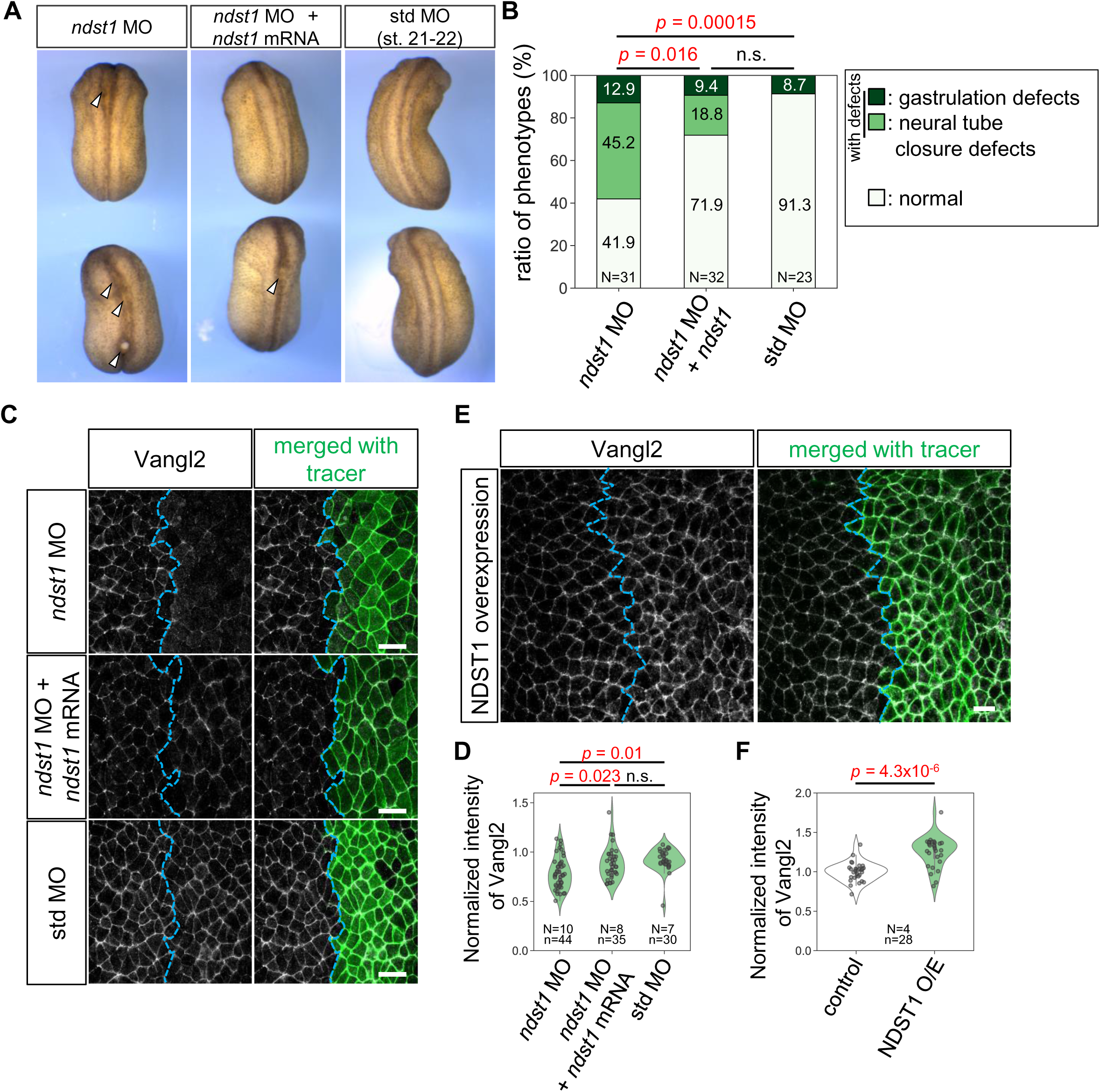
NDST1 is required for proper formation of PCP in the neural plate. (A) Morphological appearances of injected embryos (Stage 21-22). Knockdown of *ndst1* resulted in neural tube closure defects (arrowheads). MOs and *ndst1* mRNA were injected into both dorsal blastomeres of 4-cell embryos. (B) Quantification of morphological phenotypes in *ndst1* knockdown embryos (Stages 21-22). For statistical analysis, gastrulation defects and neural tube closure defects were summed as “with defects.” Statistical analysis was performed using Pearson’s Chi-squared test for pre-analysis and multiple comparison of population proportions employed Ryan’s method. Numbers of embryos (N) as indicated. (C) Requirement of *ndst1* for membrane localization of Vangl2 in the neural plate (Stage 14). MOs and *ndst1* mRNA with membrane tracer (lyn-mTagBFP2-3HA) were co-injected into the right dorsal blastomere of 4-cell embryos, targeting the future neural plate. Boundaries of tracer-negative and -positive area are indicated with dashed lines. (D) Quantification of membrane localization of Vangl2 in *ndst1* knockdown experiments (C). Statistical analysis was performed with the Shapiro-Wilk test to assess data normality, and was followed by the Kruskal-Wallis test for pre-analysis and the Steel-Dwass-Critchlow-Fligner test for multiple comparisons. Numbers of embryos (N) and numbers of sub regions (n) are as indicated. (E) NDST1 overexpression increased Vangl2 on the membrane (Stage 14). mRNAs of *ndst1* and membrane tracer (lyn-mTagBFP2-3HA) were co-injected into the right dorsal blastomere of 4-cell embryos, targeting the future neural plate. Boundaries of tracer-negative and -positive area are indicated with dashed lines. (F) Quantification of membrane localization of Vangl2 in NDST1 overexpression experiments. (E). Statistical analysis was performed with the Wilcoxon rank-sum test after the Shapiro-Wilk test to assess data normality. Numbers of embryos (N) and numbers of cell boundaries (n) are as indicated. Amounts of mRNAs/MOs: *ndst1*: 67 pg/embryo (A and B), 33 pg/embryo (C and D), 500 pg/embryo (E and F); *lyn-mTagBFP2-3ha,* 83 pg/embryo (C and D), 100 pg/embryo (E and F); *ndst1* MOs, 43 ng/embryo (A and B), 14 ng/embryo (C and D); std MO, 42 ng/embryo (A and B), 14 ng/embryo (C and E). Scale bars, 50 μm (C and D).

### All three modifications of HS chains are polarized similarly to Wnt11 and core PCP components in *Xenopus* neural plate

To gain further insight into the mechanism by which HS chains are involved in polarized Wnt11 localization and PCP formation in the neural plate, we examined spatio-temporal distribution of HS chains by whole-mount immunostaining (Figures 3A, 3B, S4C, and S4D). Surprisingly, HS chains visualized by the three antibodies exhibited clearly polarized patterns in the neural plate at Stage 14, when PCP is established (Figures 3B and S4D). At this stage, all three modification states of HS chains were significantly and highly polarized compared with wheat germ agglutinin (WGA) staining, which is used as a membrane marker (Figure S4A). Polarization of HS chains was significantly stronger at Stage 14 than Stage 12 (Figure 3C). Hence, the polarized pattern of HS chains was organized or at least enhanced within 2-3 hours, from Stage 12 to Stage 14. These results indicate that the spatial pattern of HSPGs is not static, but is dynamic during formation of the PCP.

**Figure 3.**
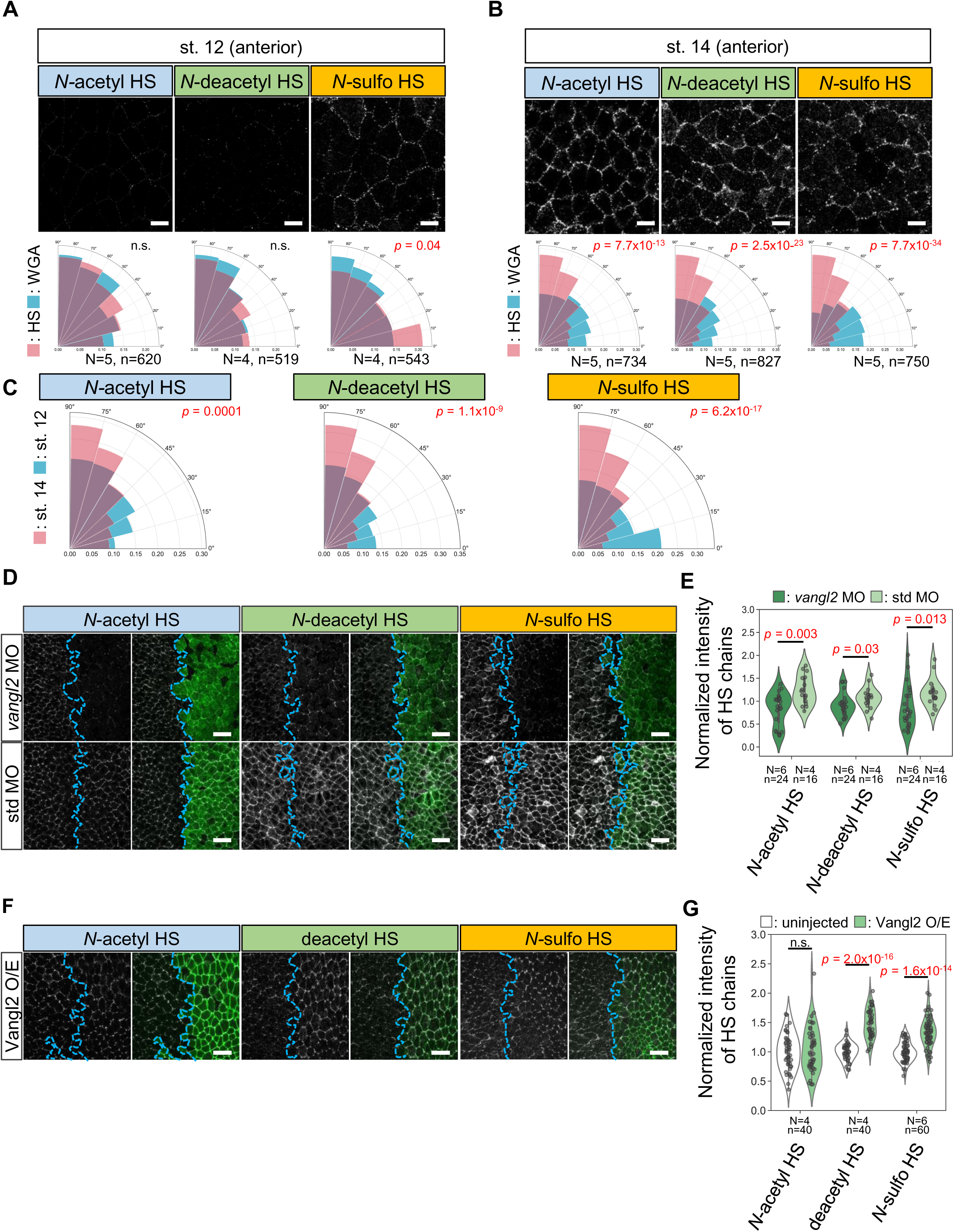
Polarized localization of HS chains in the neural plate, depends on PCP formation. (A and B) Spatio-temporal distribution of HS chains in the neural plate at Stages 12 (A) and 14 (B), corresponding to a 2-3-h interval. A relatively anterior region is presented (see also Figure 4B for the observed region). Polarization of HS chains (red rose diagram) was compared with WGA (blue rose diagram). Statistical analysis was performed with the Kuiper 2 sample test. Numbers of embryos (N) and numbers of cells (n) are as indicated. (C) Comparison of HS-chain polarization between Stages 12 and 14. Statistical analysis was performed with the Kuiper 2 sample test. (D) Effect of *vangl2* knockdown on HS chains in the neural plate (Stage 14). MOs and *vangl2* mRNA with tracer (FITC-dextran; 8.3 ng/embryo) were co-injected into the right dorsal blastomere at the 4-cell stage. Boundaries of tracer-negative and -positive area are indicated with dashed lines. (E) Quantification of membrane localization of HS chains with *vangl2* knockdown (D). For statistical analysis, the Wilcoxon rank-sum test was performed. Numbers of embryos (N) and numbers of sub regions (n) are as indicated. (F) Increase of *N*-deacetyl and *N*-sulfo HS chains on Vangl2 overexpressing cells (Stage 14). mRNA of *vangl2* and membrane tracer (mRuby2-KRas) were co-injected into the right dorsal blastomere of 4-cell embryos. (G) Quantification of membrane localization of HS chains with Vangl2 overexpression (F). For statistical analysis, the Shapiro-Wilk test was initially performed to assess data normality, and *t*-test was performed after the test of homogeneity of variance by *F*-test. Numbers of embryos (N) and numbers of cell boundaries (n) are as indicated. Amounts of mRNAs/MOs: *vangl2*: 10 pg/embryo (D and E), 100 pg/embryo (F and G); *mRuby2-kras*: 100 pg/embryo (F and G); *vangl2* MOs and std MO: 14 ng/embryo (D and E). Scale bars, 20 μm (A and B), 50 μm (D and F).

### The polarized pattern of HS chains is regulated by Vangl2 in the neural plate

The apparent coincidence between PCP establishment and polarization of HS chains prompted us to examine whether disruption of PCP affects spatial distribution of HSPGs. Thus, we knocked down *vangl2* in the right half of the neural plate using *vangl2* MO ^38,39^. We found that localization to the cell membrane of HSPGs enriched with all three modification states of HS chains was significantly reduced by the *vangl2* MO compared to the std MO (Figures 3D and 3E). Thus, Vangl2 was required for localization of HSPGs with *N-*acetyl-, *N*-deacetyl-, or *N*-sulfo-rich HS chains on cell membranes. Consistently, overexpression of Vangl2 increased localization of HSPGs with *N*-deacetyl- or *N-*sulfo-rich HS chains on cell membranes (Figures 3F and 3G). These data suggest that core PCP components are involved in localization of HSPGs in the neural plate.

### *N*-Deacetyl and *N*-sulfo HS chains can be polarized in ectopically established PCP

Localization of HS chains appears to link with PCP in the neural plate, where PCP is naturally established; however, polarization of HSPGs may be initiated by some pre-patterns or anisotropy, such as tissue tension ^37^ and/or cytoskeletons ^40^ that are independent of PCP. Thus, we next asked whether localization of HS chains is dependent on ectopic establishment of PCP. The animal cap region has only weak polarity, if any, at early neurula stages (Stages 13-14). Overexpression of Wnt11 and core PCP components (GFP-Pk3 and HA-Vangl2) in adjacent areas in this region showed that PCP can be established in an exogenous Wnt-dependent manner ^8^. Using this system (Figure 4A), we examined localization of HS chains with or without exogenous Wnt11. Without overexpressed Wnt11, GFP-Pk3, which serves as a readout of PCP, exhibited uniform distribution (Figures 4B and 4C, upper panels). In this condition, distributions of *N*-deacetyl and *N*-sulfo HS are also uniform. In contrast, *wnt11* overexpression led to clear polarization of GFP-Pk3 (localized to distal edges from the Wnt11 source) and Wnt11 itself (Figures 4B and 4C, lower panels ^15^). In this condition, *N*-deacetyl and *N-*sulfo HS exhibited polarized distributions, co-localizing with Wnt11 and GFP-Pk3 (Figures 4B and 4C, lower panels, arrowheads). This experiment indicates that the polarized distribution of HS chains can be generated upon PCP establishment without any pre-patterns of HS chains. Furthermore, co-localization of those HS chains, GFP-Pk3 and Wnt11 suggest complex formation among them. Importantly, overexpression of Wnt11 alone never caused polarization of HS chains (see Figure S6C), suggesting that establishment of PCP upon proper combination of Wnt11 and core PCP components can polarize distribution of HS chains. Indeed, correlation between Wnt11 and HS chains was significantly higher with overexpressed core PCP components (GFP-Pk3 and Vangl2) than that without (Figure 4D). Similarly, correlation between GFP-Pk3 and HS chains was significantly higher with overexpressed Wnt11 than without (Figure 4E). Overall, polarized distribution of HS chains can be generated upon PCP establishment that is driven by Wnt11, through complex formation among NDST1-modified HS chains, core PCP components, and Wnt11.

**Figure 4.**
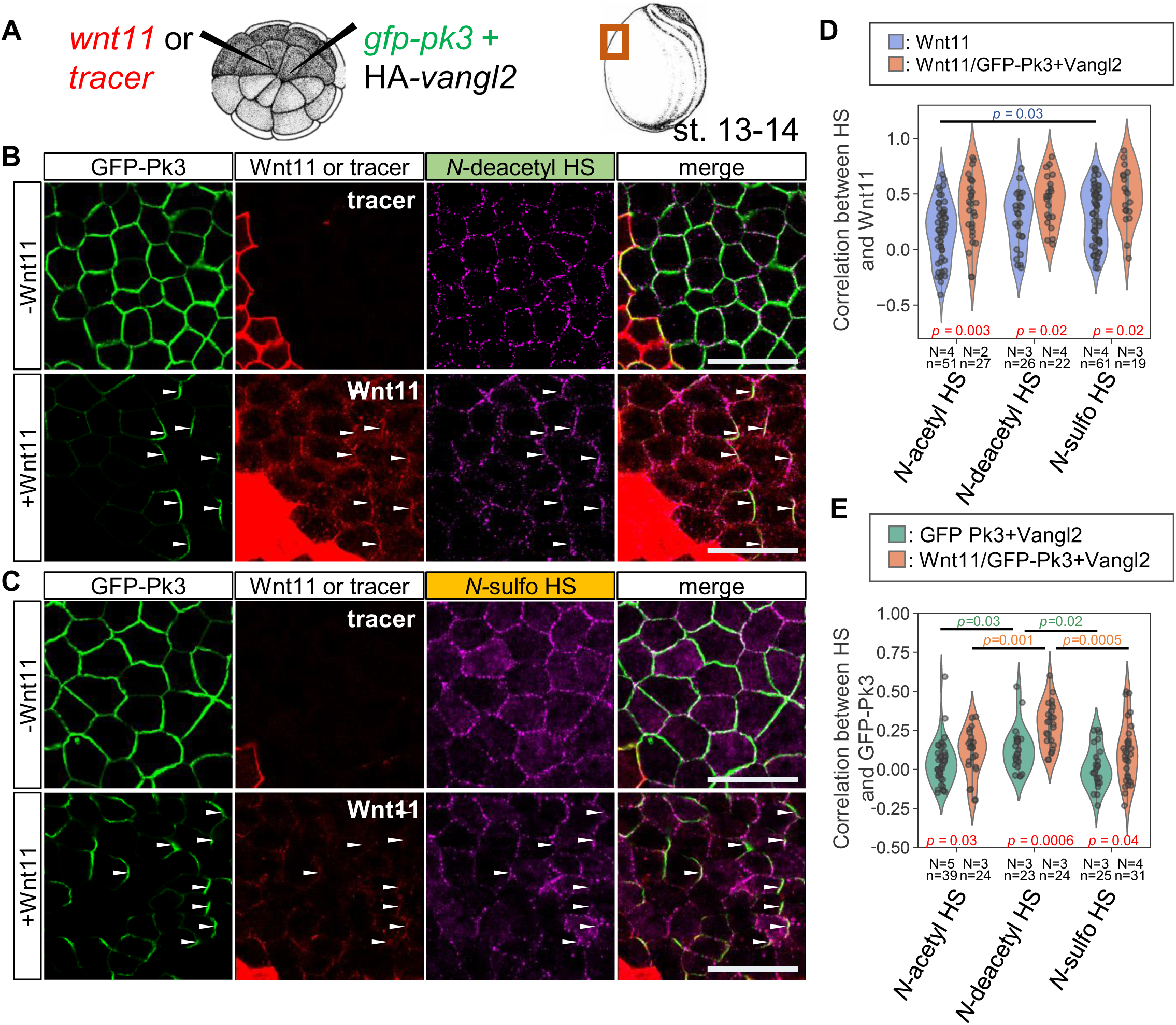
Deacetyl HS and *N-*sulfo HS can be polarized in the ectopically established PCP directed by exogenous Wnt11. (A) Schematic image of injection. mRNAs of *GFP-pk3* mixed with *ha-vangl2* and *wnt11* or tracer (mRuby2-KRas) were injected into adjacent animal-ventral blastomeres of 32-cell stage embryos. (B and C) Localization of GFP-Pk3, Wnt11, and *N*-deacetyl HS (B) or *N*-sulfo HS (C) with (lower panels) or without exogenous Wnt11 (upper panels) (Stage 14). Rabbit anti-Wnt11 antibody was generated and evaluated to visualize Wnt11 (Figure S5A). Co-localization of GFP-Pk3, Wnt11, and HS chains are indicated with arrowheads. (D and E) Quantification of correlation coefficients between HS chains and Wnt11 (D) or GFP-PK3 (E) in the ectopically established PCP or Wnt11 or core PCP components-only overexpression (see also Figures S6A and S6B). All data were performed the Shapiro-Wilk test to assess data normality. For pairwise comparison, the Wilcoxon rank-sum test was performed for *N*-sulfo HS in (D), and for *N*-acetyl and *N*-deacetyl HS in (E). For *N*-acetyl and *N*-deacetyl HS in (D) and *N*-sulfo HS in (E), a *t*-test was performed after the test of homogeneity of variance with an *F*-test. For multiple comparison in (D) and in (E, core PCP components-only overexpression), the Kruskal-Wallis test for pre-analysis and the Steel-Dwass-Critchlow-Fligner test were performed. For multiple comparison in (E, ectopically established PCP), Tukey’s HSD test was performed. For (D), numbers of embryos (N) and numbers of cell boundaries (n) are as indicated. For (E), numbers of embryos (N) and numbers of cells (n) are as indicated. Amounts of mRNAs: *GFP-pk3*, 100 pg/embryo (B - E); *ha-vangl2*, 50 pg/embryo (B and C); *vangl2*, 50 pg/embryo (D and E); *wnt11*, 250 pg/embryo (B and C); *wnt11-4ha*, 250 pg/embryo (D and E); *mRuby2-kras*, 100 pg/embryo (B - E); *mEGFP-kras*, 100 pg/embryo (D and E). Scale bars, 50 μm (B and C).

### Mechanisms underlying the association between HS chains and core PCP components in a polarized manner

To obtain mechanistic insights into the relationship between HS chains and PCP, especially association of HS chains and core PCP components in a polarized manner, we focused on Frizzled 7 (Fzd7), a receptor of Wnt11, as well as one of the core PCP components ^41^. Importantly, HSPGs act as reservoirs of Wnt ligands to potentiate Wnt signaling. For example, Glypican3 and Glypican4 interact with Fzd to enhance recruitment of Wnt ligands ^42,43^. Moreover, in canonical Wnt signaling, we have shown that *N*-sulfo HS acts as a scaffold to accumulate Fzd8, co-receptor LRP6 and Wnt8 together to form signaling complexes called signalosomes ^29^. First, co-localization of HS chains and endogenous Fzd7 in the neural plate was evaluated by whole-mount immunostaining. *N-*acetyl and *N*-deacetyl HS were slightly co-localized with Fzd7, and *N*-sulfo HS exhibited highest correlation with Fzd7 among those types of HS chains (Figures 5A and 5B), which is similar to the relationship with Wnt11 (Figures 1C and 1D). Based on this finding, we examined the requirement of *N-*sulfo HS for membrane localization of Fzd7 by *ndst1* knockdown in the neural plate. *ndst1* MO, *ndst1* MO with *ndst1* mRNA, or std MO was injected into the right half of the neural plate. Membrane localization of Fzd7 was significantly reduced by *ndst1* MO, which was partially rescued by addition of *ndst1* mRNA, showing specificity of *ndst1* MO (Figures 5C and 5E). In regulation of PCP, Wnt induces phosphorylation of Vangl2 (phospho-Vangl2) via Fzd, which is necessary for PCP formation ^44–46^. Thus, phospho-Vangl2 is a reliable readout of Wnt/PCP signaling. Similar to Fzd7, phospho-Vangl2 is highly colocalized with *N*-sulfo HS, but not with *N*-acetyl or *N*-deacetyl HS (Figure S7). We further examined the effect of *ndst1* knockdown on Wnt/PCP signaling by evaluating phospho-Vangl2. Phospho-Vangl2 was significantly reduced by *ndst1* MO compared to std MO (Figures 5D and 5F). The effect of *ndst1* MO was rescued by addition of *ndst1* mRNA, showing specificity of *ndst1* MO (Figures 5D and 5F). Similar to canonical Wnt signaling, in which *N*-sulfo HS is essential for phosphorylation of LRP6 ^29^, *N-*sulfo HS appears to be required for Wnt/PCP signaling in the *Xenopus* neural plate.

**Figure 5.**
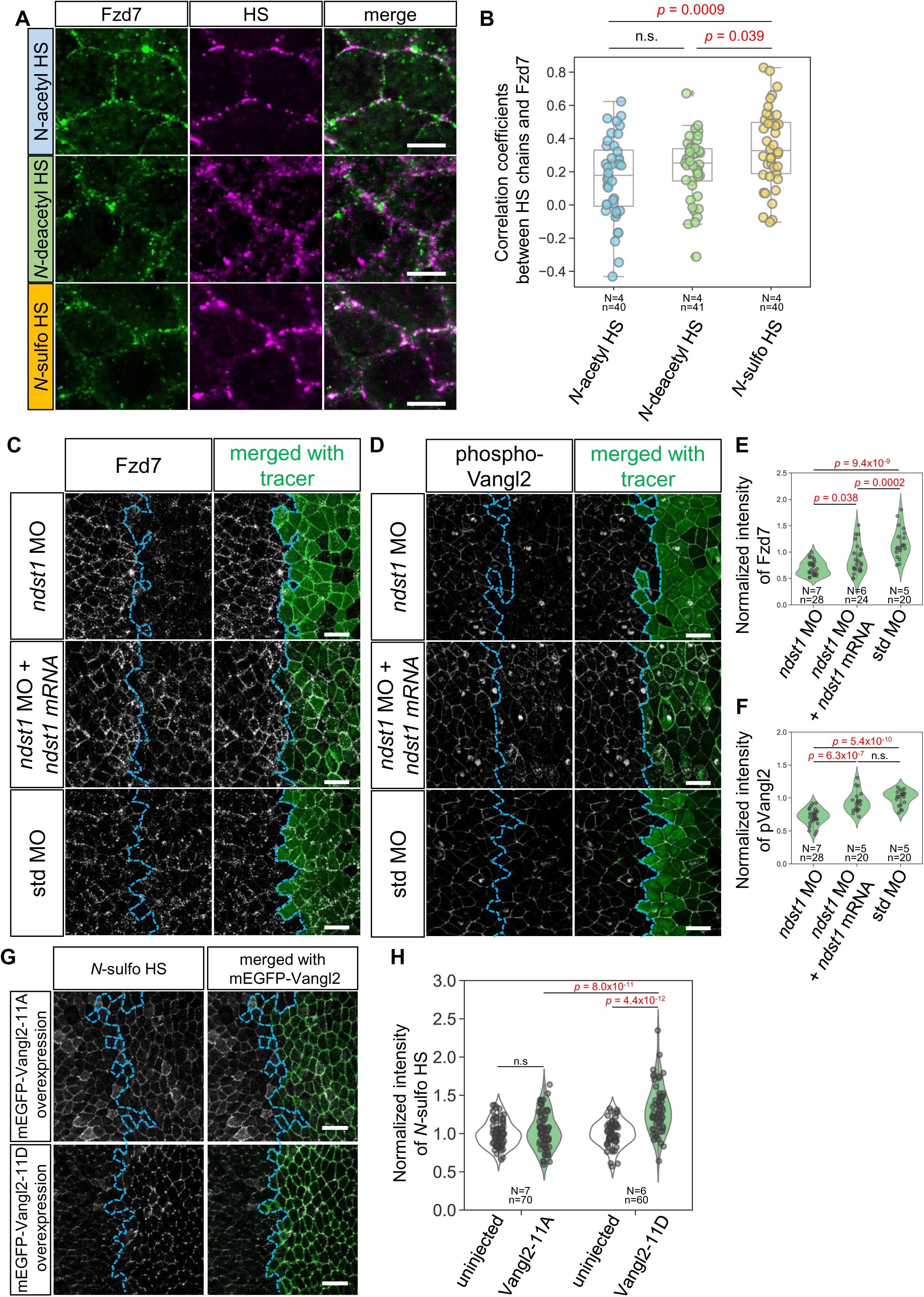
Mechanisms underlying the association between HS chains and core PCP components in a polarized manner. (A) Co-localization of endogenous Fzd7 and HS chains in the *Xenopus* neural plate (Stage 14). (B) Average of correlation coefficients of Fzd7 and HS at cell boundaries. Statistical analysis was performed with the Shapiro-Wilk test to assess data normality and was followed by Tukey’s HSD test for multiple comparisons. Numbers of embryos (N) and numbers of cell boundaries (n) are as indicated (C and D) Reduction of Fzd7 (C) and phosphorylated Vangl2 (D) from cell boundaries by *ndst1* knockdown in the neural plate (Stage 14). MOs and mRNA of *ndst1* with membrane tracer (lyn-mTagBFP2-3HA) were co-injected into a dorsal blastomere of 4-cell embryos. (E and F) Quantification of membrane localization of Fzd7 (E) and phosphorylated Vangl2 (F) in *ndst1* knockdown experiments. Statistical analysis was performed with the Shapiro-Wilk test to assess data normality and was followed by Tukey’s HSD test for multiple comparisons. Numbers of embryos (N) and numbers of sub regions (n) are as indicated. (G) Increase of *N*-sulfo HS in Vangl2-11D overexpressing cells in the neural plate (Stage 14). mRNA of *mEGFP-vangl2-11a* or *mEGFP-vangl2-11d* were injected into a dorsal blastomere of 4-cell embryos. (H) Quantification of membrane localization of *N-*sulfo HS in mEGFP-Vangl2-11A or mEGFP-Vangl2-11D overexpression experiments (G). Statistical analysis was performed with the Shapiro-Wilk test to assess data normality and was followed by Tukey’s HSD test for multiple comparisons. Numbers of embryos (N) and numbers of cell boundaries (n) are as indicated. Numbers of embryos (N) and numbers of cell boundaries (n) are as indicated. Amounts of mRNAs/MOs: *ndst1,* 33 pg/embryo (C-F); *lyn-mTagBFP2-3ha,* 83 pg/embryo (C - F); *mEGFP-vangl2-11a* or -*11d*, 500 pg/embryo (G, H); *ndst1* MOs or std MO, 14 ng/embryo (C - F), Scale bars, 10 μm (A), 50 μm (C), (D), and (G).

We have already found that Wnt11 polarizes PCP by stabilization of the asymmetric complex of core PCP components, which involves multiple loops of positive feedback regulation ^15^. We asked whether similar positive feedback regulation is involved in the association of HS chains and core PCP components. If that is the case, downstream signaling of Wnt11 would upregulate *N*-sulfo HS. Thus, we examined overexpression of phospho-mimetic mutants of Vangl2, mEGFP-Vangl2-11D, and phospho-deficient mutant, mEGFP-Vangl2-11A as a negative control ^15^. As we expected, *N*-sulfo HS was significantly increased by mEGFP-Vangl2-11D overexpression, but not by mEGFP-Vangl2-11A overexpression (Figures 5G and 5H). From these data, we suggest that *N*-sulfo HS, Wnt, Fzd7, and phosphorylation of Vangl2 are involved in positive feedback regulation to achieve polarization of all these factors in the neural plate.

## Discussion

### Dynamic pattern formation of HSPGs in association with PCP

Based primarily on intensive genetic studies, HSPGs have been considered static and uniform scaffolds on cell surfaces for graded distributions of morphogens ^17^. Genetic interactions between HSPGs and core PCP components have been reported ^20,24^, and HSPGs also exhibit genetic and physical interactions with Wnt11 ^21,22^. However, detailed cellular mechanisms in which HSPGs are involved in PCP are largely unknown. In contrast, in this study, visualization by staining with HS chain-specific antibodies demonstrated that the distribution of HSPGs changes dynamically in the neural plate, resulting in formation of a polarized pattern along the medio-lateral axis of the embryo (Figures 3A - 3C). Because similar polarized patterns of HS chains were established, depending on ectopically expressed Wnt11 and core PCP components, even in the area where PCP is not apparent in normal development, it is likely that polarization of HSPGs arises without any pre-patterns (Figures 4B and 4C). Overall, HSPGs are likely to be organized in polarized complexes with core PCP components and Wnt11 to regulate PCP, as evidenced by mutual requirements (Figures 1F, 1G, 2C, 2D, 3D, and 3E) and ectopic establishment of PCP (Figures 4B and 4C), and co-localization (Figures 4B, 4C, 5A, and S7). Thus, our findings suggest that HSPGs are dynamically organized in complexes of Wnt11 and core PCP components, resulting in the polarized pattern of HSPGs and Wnt11 ^15^. Hence, the reported genetic interaction may result from complex formation among HSPGs, Wnt11, and core PCP components. Disruption of PCP/noncanonical Wnt signaling causes several types of diseases, such as Robinow syndrome and spina bifida ^4^. Our findings may provide new insights into mechanisms of PCP-related diseases, resulting in therapeutics against these diseases.

### Assembly of signaling complexes for noncanonical Wnt/PCP signaling by HSPGs

Mechanisms underlying assembly of signaling complexes for noncanonical Wnt signaling remain unclear. In canonical Wnt signaling, signaling components, such as Fzd, LRP5/6, and Dishevelled, are clustered to assemble signaling complexes called signalosomes, leading to phosphorylation of LRP6, a crucial step in activation of the canonical pathway ^47–49^. We reported that clusters of *N*-sulfo-rich HSPGs serve as scaffolds of signalosome assembly by accumulating Wnt8, Fzd8, and LRP6 ^29^. Similarly, we suppose that *N*-sulfo-rich HSPG clusters also serve as scaffolds to assemble signalosome-like complexes for noncanonical Wnt signaling because *N-* sulfo HS and endogenous Fzd7 are specifically co-localized (Figures 5A and 5B), and *ndst1* is required for membrane localization of Wnt11 (Figures 1F and 1G) and Fzd7 (Figures 5C and 5E) and downstream phosphorylation of Vangl2 in the neural plate (Figures 5D and 5F).

In *Drosophila*, although actual involvement of Wnt ligands in regulation of PCP is debatable^10,11^, signalosome-like assembly of core PCP components, including phosphorylated Vang/Stbm, are reported ^50,51^. Association of HSPGs and core PCP components is also suggested in *Drosophila*, evidenced by reduction of Glypican-type core proteins of HSPGs in mutant clones or knockdown of *pk* or *vang* ^52^. Our data further suggest an association between HS chains and core PCP components (Figures 2C - 2F and 3D - 3G). Of note, the signalosome-like assembly in *Drosophila* occurs over a wide range of stoichiometries of core PCP components ^50^. HS chains may contribute to allow such a wide compositional range, because they are highly repetitive polymers of disaccharide units, which generally have a high capacity and low or moderate affinity for protein binding.

### Limitations of this study

We found some differences between NDST1-modified HS chains, *N*-deacetyl, and *N*-sulfo HS in co-localization with Wnt11 (Figures 1C and 1D) and Fzd7 (Figures 5A and 5B). However, because NDST1 has both *N*-deacetylase and *N*-sulfotransferase activities, it is difficult to segregate specific functions of these modified states. Further analysis will be required to clarify possibly distinct roles of deacetyl and *N*-sulfo HS in PCP, for example, by specific manipulations of these modifications.

In addition, core proteins of HSPGs responsible for the polarized pattern of HS chains are not determined in this study. Glypican4 and Syndecan4 are involved in PCP ^21–24^. Although, we did not examine core proteins in this study, relationships between core proteins and HS modifications in regulation of PCP need to be addressed in future work, because core proteins show specificity for HS modifications ^29^.

## Methods

### *Xenopus* embryo manipulation and microinjection

All experiments using *Xenopus laevis* were approved by the Institutional Animal Care and Use Committee, National Institutes of Natural Sciences or the Animal Experimentation Committee, Kyoto University. Manipulation of *Xenopus laevis* and microinjection experiments were performed according to standard methods ^53^. Briefly, unfertilized *Xenopus* eggs were obtained by injection of gonadotropin (ASKA Pharmaceutical), and eggs were fertilized with testis homogenates. After fertilization, eggs were de-jellied in 4% cysteine solution (pH7.8) and incubated in 1/10x Steinberg’s solution at 14-20 °C. Embryos were staged according to Nieuwkoop and Faber ^34^. For mRNA microinjection, mRNAs were synthesized from plasmid DNA with an mMessage mMachine SP6 kit (Invitrogen) and purified with an RNeasy Micro kit (QIAGEN). Microinjections were performed in early (4-32 cell) embryo. Amounts of injected mRNA are described in figure legends. For Morpholino oligo (MO) injection, 6-9 ng of FITC-dextran (Molecular Probes, D1820), or mRNA encoding mEGFP-KRas, mRuby2-KRas or lyn-mTagBFP2-3ha were used for cell tracing. Amounts of injected mRNAs/MOs are described in the figure legends.

### Plasmids used for mRNA synthesis

We used the following plasmids for templates of mRNA synthesis: pCSf107-*ndst1* ^17^; *vangl2*, *wnt11*, *wnt11-mTagBFP2, mEGFP-vangl2-11d*, *mEGFP-vangl2-11d, mRuby2-kras*, *mEGFP-kras*, and *lyn-mTagBFP2-3ha* ^15^; pXT7-*gfp-pk3* and pCS2+*ha-vangl2* ^8,54^.

### Morpholino oligos (MOs)

Because *X. laevis* is an allotetraploid, MOs were designed to target transcripts from both homoeologs and were mixed at the same concentration. ndst1, AGGAATGGCACAAGCTCACAAATGC and AGGAGTGGCACAAGCTCACAAATGC ^29^; vangl2, GAGTACCGGCTTTTGTGGCGATCCA ^15,38^ and CGTTGGCGGATTTGGGTCCCCCCGA ^39^ std MO (CCTCTTACCTCAGTTACAATTTATA) was used as a negative control ^29^.

### Fluorescent image acquisition

Fluorescent images were acquired mainly with a laser-scanning confocal microscope (Leica TSC SP8 system, objective: HC PL APO2 x40/NA1.10 W CORR water immersion) and partly with a spinning disc confocal microscope (Andor Dragonfly 200 combined with an Olympus IX83, objective: UPLXAPO20X x20/NA0.8).

### Live-imaging

For live-imaging, embryos were mounted on 35-mm glass-based dishes (Iwaki, 3910-035) using a silicone chamber made in-house with holes 1.8 mm in diameter.

### Immunostaining

*Xenopus* embryos were fixed with MEMFA (0.1M MOPS, pH 7.4, 2mM EGTA, 1mM MgSO4, 3.7% formaldehyde) and dehydrated with EtOH to improve HS chain staining, except for Figures 5C and 5D, because EtOH dehydration is not good for staining Fzd7 or pVangl2. Fixed embryos were rehydrated with TBT (1x TBS, 0.2% BSA, 0.1% Triton X-100). For Vangl2 staining, antigen retrieval was performed at 95 °C for 20 min in citrate-based Antigen Unmasking Solution (H-3300-250, Vector Laboratory) after rehydration. Blocking was performed with TBTS (TBS with 10% heat-immobilized FBS [70 °C, 40 min]). Primary antibodies and their dilutions were as follows: anti-Wnt11 (rabbit polyclonal IgG, in-house preparation, 1/2000, or mouse monoclonal IgG2b, in-house preparation, 1/10 ^15^), anti-Vangl2 (HPA027043, Sigma, rabbit polyclonal IgG, 1/200), anti-Fzd7 (ab64636, Abcam, rabbit polyclonal IgG, 1/200), anti-Phospho-Vangl2-T78/S79/S82 (AP1206, ABclonal, rabbit polyclonal IgG, 1/1000), anti-HA (11867423001, Roche, 3F10, rat monoclonal IgG1, Roche, 1/2000), anti-C-Cadherin (6B6, Developmental Studies Hybridoma Bank, mouse monoclonal IgG1, 1/50), anti-ZO1 (33-9100, Invitrogen, ZO1-1A12, mouse monoclonal IgG1, 1/200) anti-*N*-acetyl HS (in-house preparation, NAH46, mouse monoclonal IgM, 1/50 ^29^), anti-*N*-sulfo HS (in-house preparation, HepSS-1, mouse monoclonal IgM, 1/400 ^29^), and anti-*N*-deacetyl HS (370730-1, Amsbio, JM-403, mouse monoclonal IgM, 1/200). Primary antibodies were diluted with Can Get Signal immunostain Immunoreaction Enhancer Solution A (NKB-501, TOYOBO, for Figure 1A) or B (NKB-601, TOYOBO, for Figures 1F, 5C, and 5D) or TBTS, unless specified. Secondary antibodies and their dilutions were as follows: goat polyclonal anti-mouse IgM mu chain Alexa Fluor 555 (A21426, Molecular Probes, 1/500), goat polyclonal anti-mouse IgM mu chain Alexa Fluor 647 (ab150123, Abcam, 1/500), goat polyclonal anti-mouse IgG1 (γ1) Alexa Fluor 488 (A21121, Molecular Probes, 1/500), goat polyclonal anti-mouse IgG Alexa Fluor 488 (A11001, Molecular Probes, 1/500), goat polyclonal anti-mouse IgG Alexa Fluor 647 (A21235, Molecular Probes, 1/500), goat polyclonal anti-rabbit IgG Alexa Fluor 488 (A11008, Molecular Probes, 1/500), goat polyclonal anti-rabbit IgG Alexa Fluor 555 (A21429, Molecular Probes, 1/500), goat polyclonal anti-rat IgG Alexa Fluor 555 (A21434, Molecular Probes, 1/500). Secondary antibodies were diluted with TBTS. Embryos were incubated in antibody solution overnight at 4 °C and washed with TBT 6 x 30 min after incubation with primary or secondary antibodies. Wheat germ agglutinin (WGA)-conjugated with Alexa Fluor 647 (W32466, Molecular Probes) was mixed in secondary antibody solution at dilution of 1/2000.

For double staining of *N*-deacetyl HS and *N*-sulfo HS chains, anti-*N-*sulfo HS (270426, HepSS-1, Seikagaku) was directly labeled with Alexa Fluor 488 5-SDP ester (A30052, Molecular Probes). Briefly, the antibody solution (1.0 mg/mL in PBS) was mixed 1/40 with reactive dye (10 mg/mL in DMSO) for conjugation reaction. The mixture was then incubated for 1.5 h at room temperature with shading. For purification of labeled antibody, unreacted dye was removed by gel filtration using Bio-Spin 6 Tris Columns (#732-6227; Bio-Rad). For staining of *N*-deacetyl HS chain (JM-403), primary antibody (JM-403) and secondary antibody (Cy3 AffiniPure Fab Fragment Goat Anti-Mouse IgM (µ-chain-specific) (115-167-020, Jackson ImmunoResearch)) were diluted x 1/200 with TBTS. Staining procedures were as described above. After staining for *N*-deacetyl HS chains, embryos were incubated with direct-labeled HepSS-1 (x 1/5 dilution in TBTS) overnight at 4 °C and washed with TBT three times for 1.5 hr in total.

### Statistical analyses

Sample sizes were empirically determined. The Shapiro-Wilk test was used to assess normality of datasets. Parametric tests were used for datasets with normal distributions. Otherwise, nonparametric tests were used as follows. For parametric tests of single pairs of data, Student’s or Welch’s *t*-test (two-sided) was performed after comparing variances of a set of data by *F*-test. For nonparametric testing of a single pair of data, the Wilcoxon rank-sum test was performed. Tukey’s honestly significant difference (HSD) test was performed for parametric multiple comparisons, and the Kruskal-Wallis test followed by the Steel-Dwass-Critchlow-Fligner test was performed for nonparametric multiple comparisons. For comparison of proportions in Figure 2B, Pearson’s Chi-squared test followed by Ryan’s multiple comparison of population proportions was performed. For pairwise comparisons of polarity, Kuiper 2 sample test (a circular analog of the Kolmogorov–Smirnov test) was used ^37^.

### Image analyses

All measurements for calculating signal intensity were performed with Fiji. Because basal signal intensity varied among embryos, normalized intensities of region of interest (ROI) were calculated by dividing intensities of ROIs by averaged intensity of the uninjected side for each embryo. For correlation between Wnt11-4HA and HS chains (Figures 1C and 1D), line plots were performed by tracing cell boundaries from Wnt11-4HA expressing cells with a 5-pixel width. For correlation between *N*-deacetyl and *N*-sulfo HS chains in the neural plate (Figure S1B - S1D), line plots were performed by tracing cell boundaries along medio lateral cell boundaries with a 10-pixel width. For correlation between HS chains and Wnt11 (Figure 4D), line plots were performed by tracing cell boundaries with a 3-pixel width. For correlation between HS chains and core PCP components (Figure 4E), line plots were performed by tracing around cells with a 3-pixel cell width.

For correlation between Fzd7 and HS chains in the neural plate (Figures 5A and 5B), signal intensities along medio-lateral cell boundaries were measured with a 10-pixel cell width. For intensity of endogenous Wnt11 in *ndst1* knockdown experiment (Figures 1F and 1G) and Vangl2 in NDST1 overexpression experiment (Figures 2E and 2F), signal intensities along medio-lateral cell boundaries were measured with a 5-pixel width. For Vangl2, Fzd7, and phosphorylated Vangl2 (Figures 2C, 2D, and 5C - 5F) in *ndst1* knockdown experiments, and for HS chains in *vangl2* knockdown experiments (Figures 3D and 3E) mask images for cell segmentation were created with “Cellpose” (with the model of “Cyto2”) ^55^, using C-Cadherin staining for Figures 2C, 2D, and 5C to 5F, and using WGA for Figures 3D and 3E. The cell-segmentation mask was corrected with “TissueAnalyzer” in the Fiji plugin ^56^. 4 or 5 square regions in the injected side and the uninjected side were used as ROIs in each image. For quantification of signal intensity at cell boundaries, an intersection image of the cell segmentation mask and staining images were was generated, using “AND” operation of the “Image Calculator” plugin in Fiji. In Figures 3D and 3E, the background was subtracted with a rolling ball radius of 60 pixels. For quantification of polarity of HS chains and WGA (Figures 3A - 3C and S4A, S4C and S4D), cell-segmentation mask images were made from ZO-1 staining images using “TissueAnalyzer”. Quantification of polarity was performed with “QuantifyPolarity” ^57^.

## Acknowledgements

We thank Sergei Y. Sokol for pXT7-gfp-pk3 and pCS2+ha-vangl2, Seikagaku Corp. for NAH46 antibody, and Steven D. Aird for editing and proofreading. This work was supported in part by following programs: KAKENHI (22H02637, 23H04930 to YM; 21H02498, 22H05642, 23H04930 to ST; 22J12262 to MS), JST-PRESTO (JPMJPR194B to YM). MS was a JSPS Research Fellow.

## Author contributions

The project was conceived by MS and YM with inputs from ST. YM initiated the project with preliminary analyses. MS performed almost all preliminary and formal analyses. MS and YM designed experiments. RT generated recombinant Wnt11 protein and the rabbit anti-Wnt11 antibody. TK and MM generated the mouse monoclonal anti-Wnt11. MS wrote the first draft of the paper, and ST and YM edited the manuscript. YM supervised the project with co-supervision of ST.

## Supplementary Figure Legends

**Figure S1.**
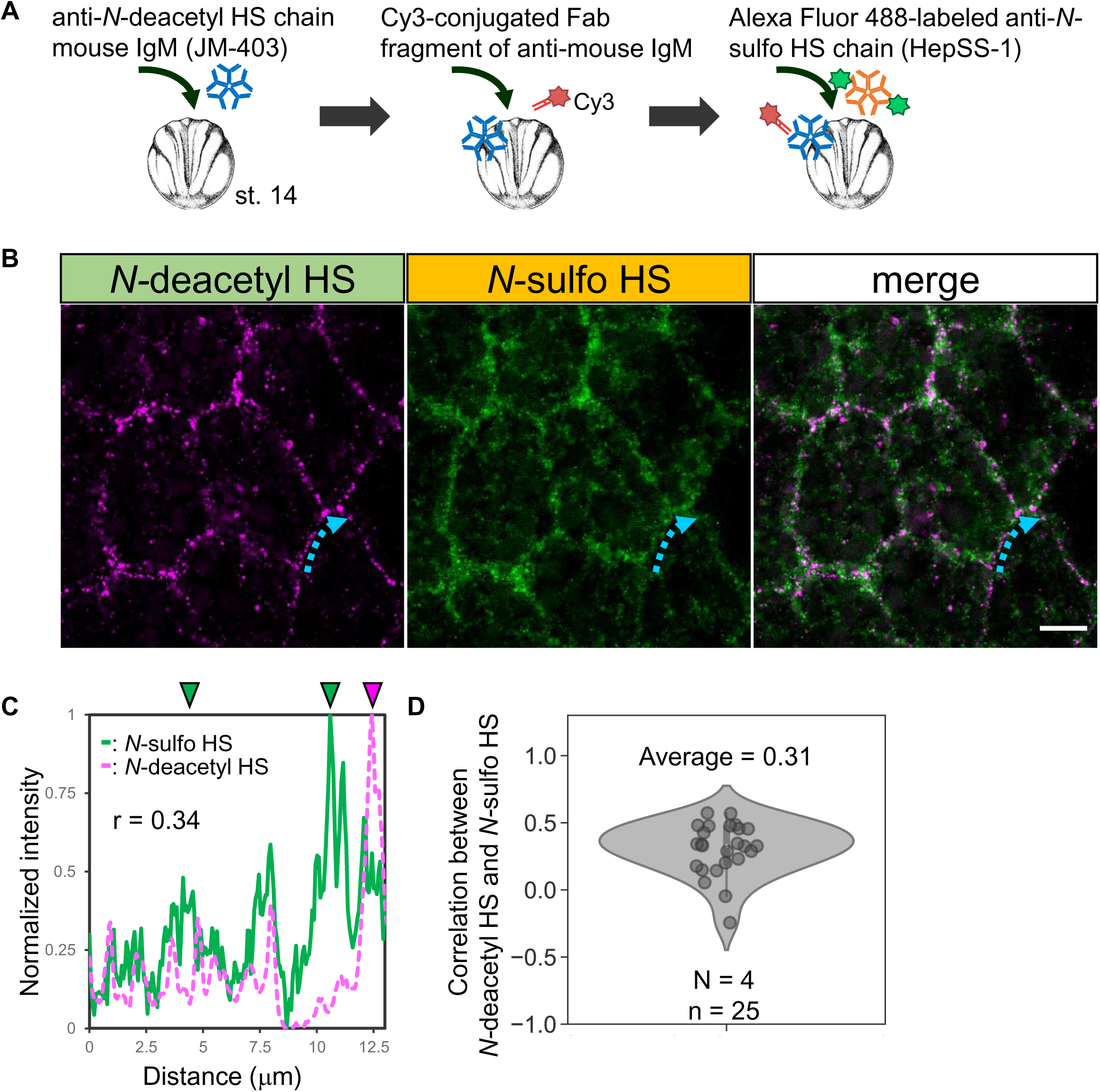
*N-*deacetyl HS and *N-*sulfo HS are in part separately clustered on cell membranes in the neural plate. (A) Strategy for double staining of *N*-deacetyl and *N*-sulfo HS chains in the neural plate. Stage 14 embryos were stained with anti-*N-*deacetyl HS (JM-403, mouse IgM) and a Cy3-conjugated monovalent Fab fragment of anti-mouse IgM to avoid cross-reaction. Then, embryos were stained with anti-*N-*sulfo HS (HepSS-1, mouse IgM) conjugated with Alexa Fluor 488. (B) Localization of *N*-deacetyl and *N*-sulfo HS in the neural plate (stage 14). The cyan dotted arrow indicates a cell boundary for the line plot of *N*-deacetyl and *N*-sulfo HS intensities in (C). (C) Comparison of localization of *N*-deacetyl and *N*-sulfo HS chains. The magenta dotted line indicates the normalized intensity of *N-*deacetyl HS staining and the green line indicates the normalized intensity of *N-*sulfo HS staining along the cell boundary shown in (B). Magenta and green arrowheads indicate positions where *N*-deacetyl and *N*-sulfo HS are highly accumulated, respectively. Fluorescent intensities were normalized with Min-Max normalization in the plot. (D) Correlation coefficients between localization of *N*-deacetyl HS and *N*-sulfo HS in the neural plate. Numbers of embryos (N) and numbers of cell boundaries (n) are as indicated. Scale bar, 10μm.

**Figure S2.**
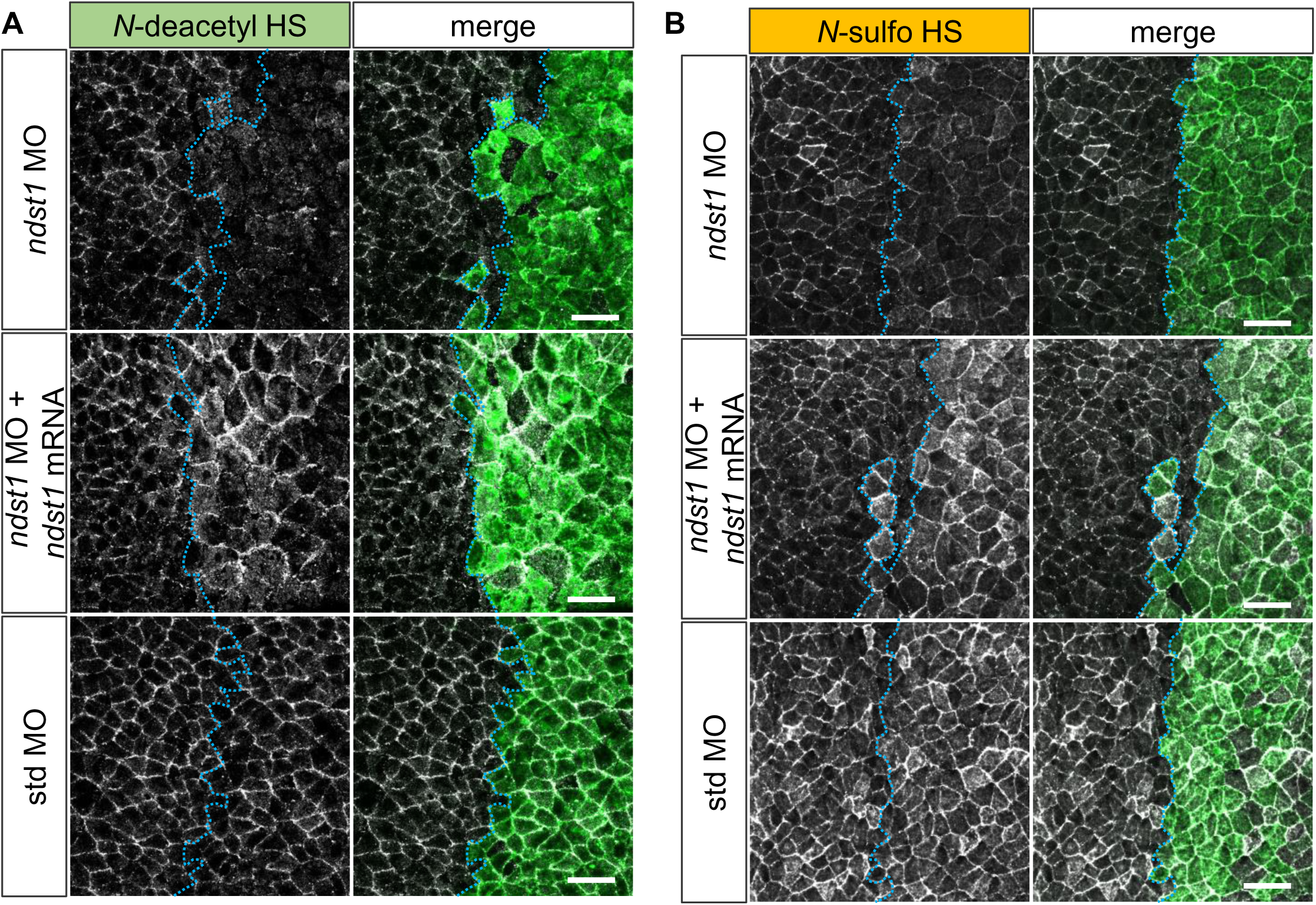
*ndst1* MO reduced *N*-deacetyl and *N-*sulfo HS on cell membranes. (A and B) *ndst1* MO reduced *N*-deacetyl HS and *N-*sulfo HS in the neural plate (Stage 14), which can be rescued by addition of *ndst1* mRNA. MOs and mRNA of *ndst1* with tracer (FITC-dextran; 8.3 ng/embryo) were co-injected into the right dorsal blastomere of 4-cell embryos. Amounts of mRNAs/MOs: *ndst1*, 13 pg/embryo; *ndst1* MO or std MO 14 ng/embryo. Scale bars, 50 μm.

**Figure S3.**
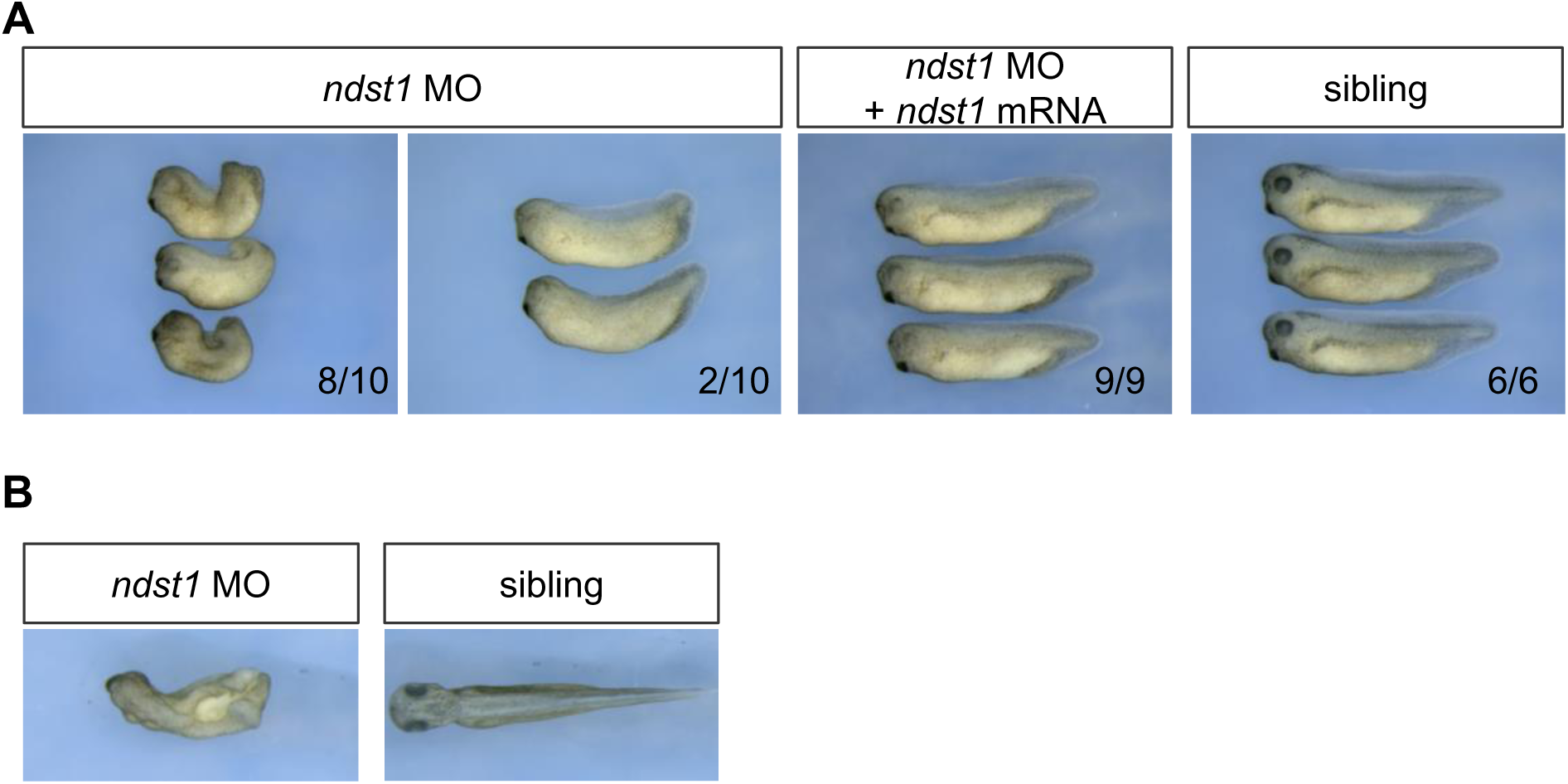
Morphological phenotypes in *ndst1* knockdown embryos. (A) Lateral view of *ndst1* MO-injected embryos in tailbud stages (Stages 35/36). MOs and mRNA of *ndst1* with tracer (FITC-dextran; 6.6 ng/embryo) were co-injected into both dorsal blastomeres of 4-cell embryos. Note that the efficiency of *ndst1* MO to cause morphological phenotype tends to vary among experiments. In this case, the effect of the MO appears more severe than in Figures 2A and 2B. (B) Dorsal view of *ndst1* MO-injected embryos in tailbud stages (Stages 35/36). Amounts of mRNAs/MOs: *ndst1*, 26 pg/embryo; *ndst1* MO or std MO 28 ng/embryo.

**Figure S4.**
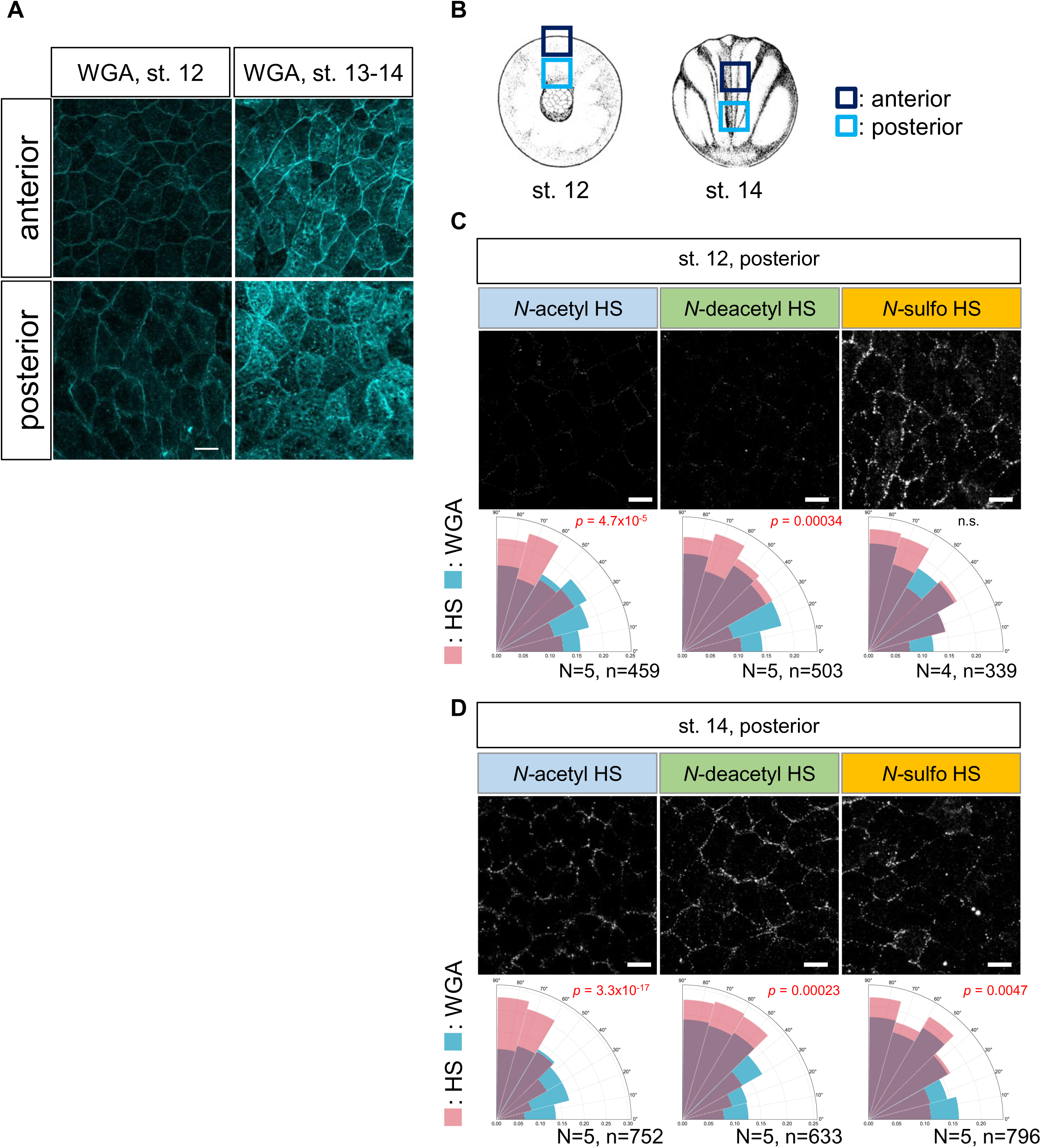
Polarized distribution of HS chains in the *Xenopus* neural plate. (A) WGA staining in the neural plate, as a membrane marker. (B) Observed areas for the distribution of HS chains (for Figures 3A, 3B, S4A, S4C, and S4D). (C and D) Distribution of HS chains in the posterior region. Polarity of HS chains and WGA staining as an internal control were quantified. Statistical analysis was performed with the Kuiper two-sample test. Numbers of embryos (N) and numbers of cells (n) are as indicated. Scale bars, 20 μm.

**Figure S5.**
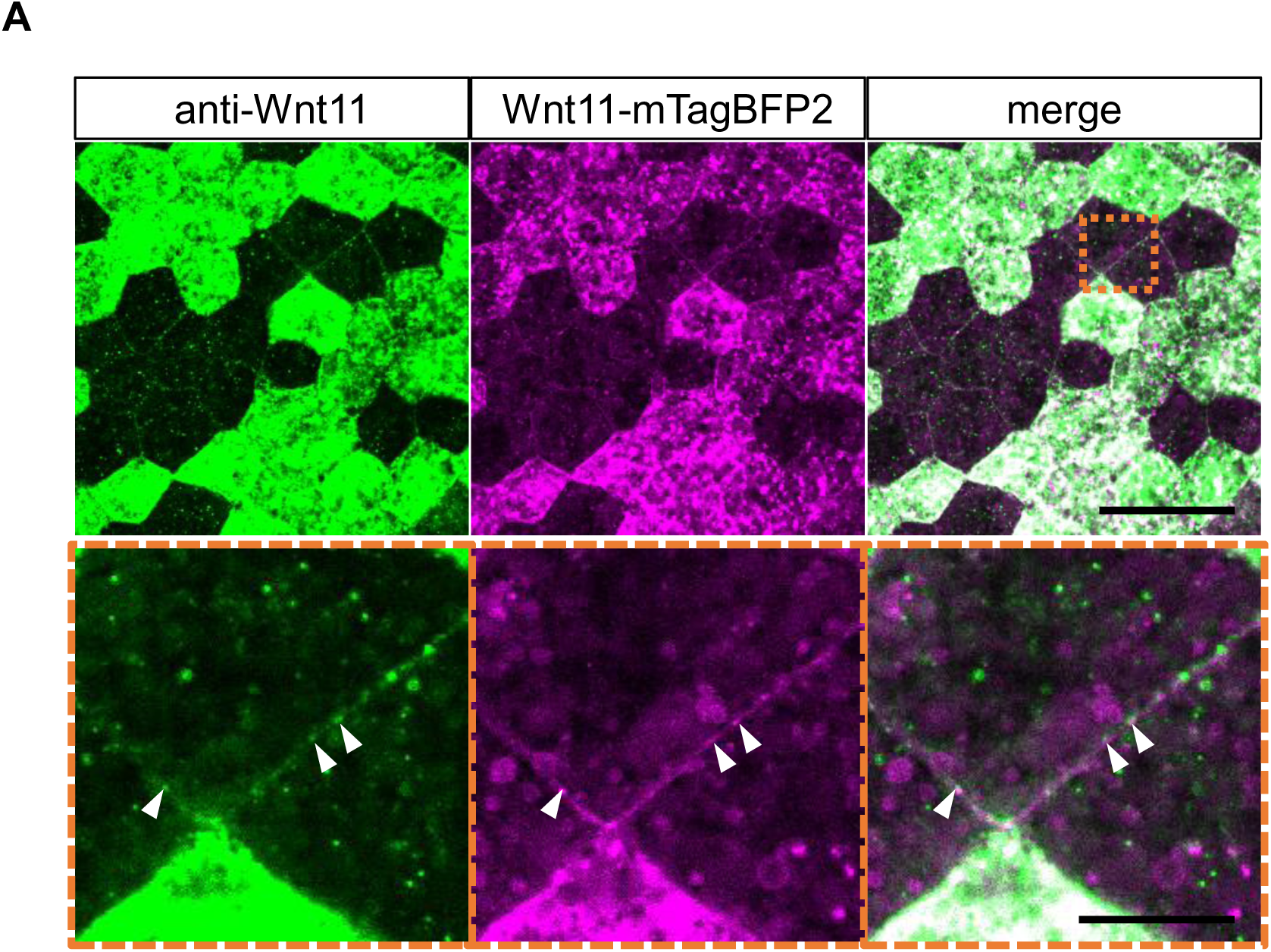
Validation of rabbit anti-Wnt11 antibody (A) Co-localization of overexpressed Wnt11-mTagBFP2 and anti-Wnt11 staining (Stage 14). *Wnt11-mTagBFP2* mRNA was injected into a ventral blastomere at the 8-cell stage. Co-localization of micro patterns were observed (lower panel with white arrow heads). Amount of mRNA: *wnt11-mTagBFP2*, 500 pg/embryo. Scale bars, 50 μm (upper panels), 10 μm (lower panels).

**Figure S6.**
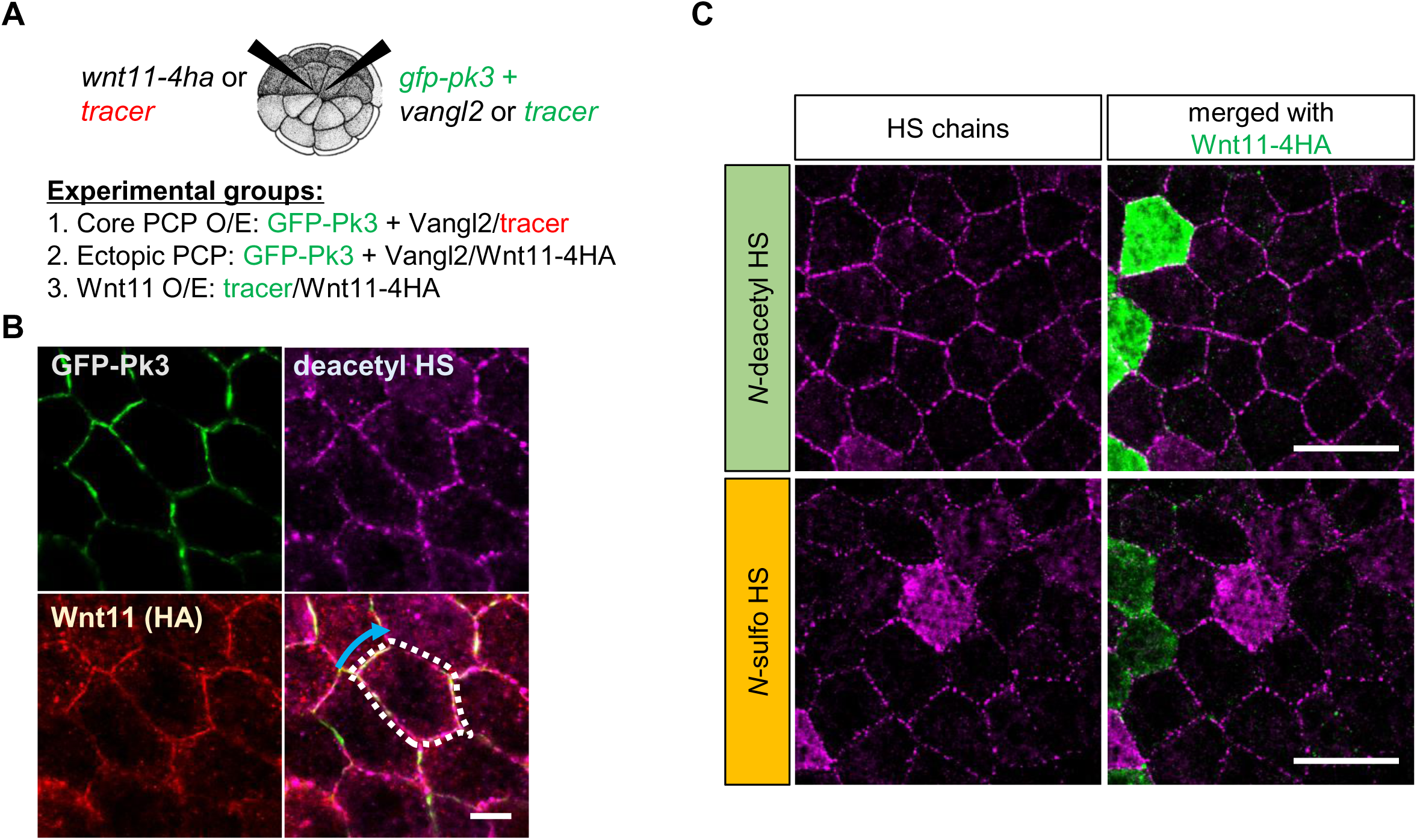
Quantification strategy of HS chains dynamics in PCP (A) Schematic image of microinjection. mRNAs of *GFP-pk3* mixed with *ha-vangl2* or tracer (mEGFP-kRas) and *wnt11* or tracer (mRuby2-kRas) were injected into adjacent ventral-animal blastomeres of 32-cell embryos. (A) Methods to measure correlation between HS chains and GFP-Pk3 or Wnt11-4HA in each experimental group (Figure S6A). For correlation between HS chains and GFP-Pk3, intensities were measured along lines surrounding cells (as exemplified with white dotted lines) with a 3-pixel width. For correlation between HS chains and Wnt11-4HA, cell edges with strong Wnt11-4HA accumulation (as exemplified with blue arrow) were measured with a 3-pixel width. (A) Overexpression of Wnt11 alone did not polarize HS chains in animal cap region (Stage 14). Amounts of mRNAs: *GFP-pk3*, 100 pg/embryo (B - E); *vangl2*, 50 pg/embryo (B and C); *wnt11-4ha*, 250 pg/embryo (D and E); *mRuby2-kras*, 100 pg/embryo (B - E); *mEGFP-kras*, 100 pg/embryo (D and E). Scale bars, 10 μm (B), 50 μm (C).

**Figure S7.**
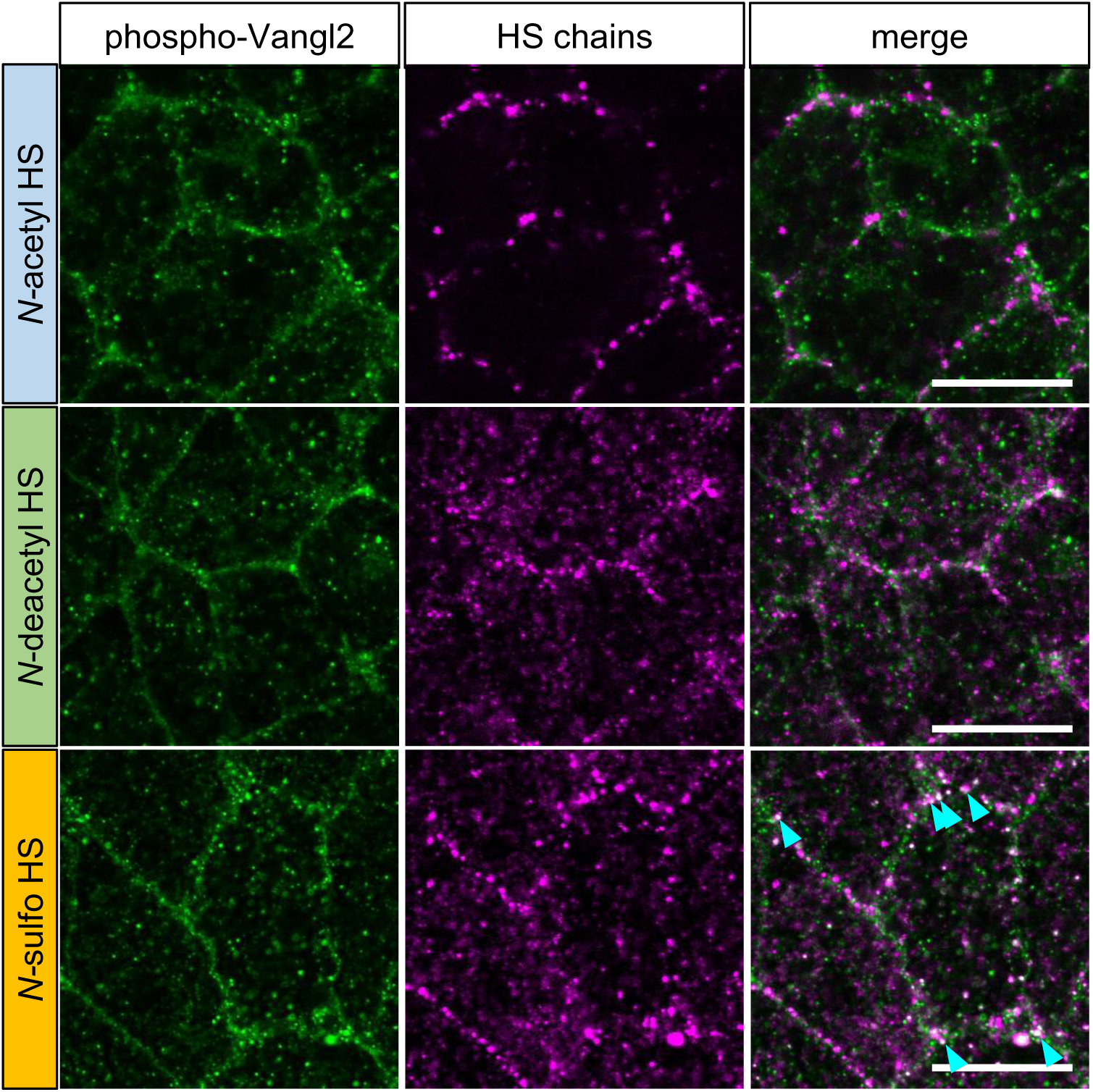
Co-localization of HS chains and phosphorylated Vangl2 in the neural plate. (A) Co-localization of HS chains and phospho-Vangl2 (pVangl2) in the neural plate (Stage 14). Cyan arrow heads indicate co-localization of *N*-sulfo HS and phosphorylated Vangl2. Scale bars, 20 μm (A).

